# Kaiso reads methylated CpGs at nucleosome entry/exit and displaces the H3 tail

**DOI:** 10.64898/2026.05.10.724166

**Authors:** Fabiana C. Malaga Gadea, Evgenia N. Nikolova

## Abstract

The zinc finger transcription factor Kaiso recognizes methylated CpG dinucleotides at silenced promoters and imprinted loci, but how it engages methylated DNA within the nucleosome remains unclear. To address this, we developed a DNMT1-based strategy for preparing site-specifically methylated nucleosomes with defined position and methylation state of the Kaiso recognition motif. Electrophoretic mobility shift assays show that Kaiso binds methylated nucleosomes with strong positional preference, with high-affinity engagement at the entry/exit site (SHL 6.5; K_d_ ≈ 100 nM), reduced affinity at SHL 5.5 (K_d_ ≈ 170 nM), and no methylation-dependent enhancement at dyad-proximal positions. Hemi- and fully methylated substrates bind Kaiso comparably at SHL 6.5, and the E535A mutation, which disrupts a key methyl-CpG contact, reduces binding in a methylation- and position-dependent manner. Solution NMR titrations of ^15^N-labeled H3 nucleosomes reveal that Kaiso binding perturbs a discrete set of H3 N-terminal tail residues, with chemical shifts trending toward free-peptide values, indicating release of the tail from its nucleosomal DNA contacts. This pattern closely resembles that produced by the pioneer factor Sox2 at the same nucleosomal region, suggesting H3 tail displacement is a general consequence of factor engagement at the nucleosome edge, independent of DNA-recognition mode. These results establish Kaiso as an active reader of methylated nucleosomal DNA that may prime local chromatin by exposing the H3 tail.

## Introduction

Gene regulation in eukaryotes requires the coordinated interpretation of multiple layers of epigenetic information, including DNA methylation,(1-3) histone post-translational modifications (PTMs),(4-6) and chromatin architecture.(7, 8) A central challenge is understanding how transcription factors (TFs) integrate these signals to achieve precise, context-dependent control of gene expression.(9-13) Kaiso (ZBTB33) is a member of the BTB/POZ (broad-complex, tramtrack, bric-à-brac/poxvirus and zinc finger) subfamily of TFs that recognizes cognate DNA sequences through a C-terminal zinc finger domain, while its N-terminal BTB/POZ domain mediates protein-protein interactions.(14, 15) What distinguishes Kaiso from other BTB/POZ family members is its unusual capacity to recognize DNA through two chemically distinct modes: a methyl-CpG-dependent mode targeting two consecutive symmetrically methylated CpG (mCpG) dinucleotides, and a sequence-specific mode that targets the unmethylated consensus TCCTGCNA motif, known as the Kaiso binding site (KBS).(15, 16) This dual specificity places Kaiso at the intersection of genetic and epigenetic readout, allowing this TF to integrate both DNA sequence and methylation status into transcriptional output.

Structural studies have revealed the molecular basis for this dual specificity.(15, 17, 18) Crystal structures of the Kaiso zinc finger domain bound to KBS and to a symmetrically methylated DNA derived from the E-cadherin promoter showed that recognition of specific bases in the major groove is achieved through both classical and methyl CH…O hydrogen bonds, while residues in a C-terminal extension bind in the opposing minor groove and are required for high-affinity binding.(15, 17) Subsequent work clarified that, despite engaging the same DNA-binding surface, the two recognition modes operate through fundamentally different mechanisms at the level of protein conformational dynamics.(17, 18) A key glutamate residue, E535, adopts distinct conformations in the two complexes: with methylated DNA, multiple direct contacts between E535 and the 5′ mCpG dominate binding affinity, whereas with KBS, E535 contributes to an indirect readout of the flanking sequence, relying on tyrosine-DNA interactions to stabilize an optimal DNA conformation.(17, 18) Together, these findings established that Kaiso exploits conformational plasticity to interpret two chemically distinct DNA signals using a single zinc-finger scaffold.

Beyond DNA recognition, Kaiso has been functionally linked to the recruitment of chromatin-modifying machinery. Kaiso is a component of the human N-CoR complex, and *in vitro* the Kaiso/N-CoR complex binds specific CpG-rich sequences in a methylation-dependent manner.(19) In vivo, Kaiso recruits the N-CoR co-repressor complex to methylated target promoters, including MTA2, where it drives histone hypoacetylation, H3K9 methylation, and the formation of repressive chromatin structures.(19, 20) In addition to its repressive functions, Kaiso regulates developmental gene expression programs through its interaction with p120-catenin,(14) which can relieve Kaiso-mediated repression of canonical Wnt target genes including c-Myc, Cyclin D1, and Wnt11.(21-26) This positions Kaiso at the interface of cell adhesion signaling and epigenetic gene regulation.

Despite this progress, how Kaiso operates in native chromatin remains unresolved. Genome-wide ChIP-seq has revealed a striking discrepancy between the *in vitro* preference for methylated DNA and the *in vivo* Kaiso binding. Rather than occupying methylated CpGs at repressed promoters, Kaiso is enriched at highly active, acetylated promoters, with nucleosome occupancy proposed to restrict access to potential binding sites in a cell type-specific manner.(27) More recent ENCODE analyses showed that Kaiso does bind methylated CpGs, but predominantly within compacted, DNase-inaccessible domains carrying the H3K4me1 signature in the absence of other active or repressive marks – a “primed” heterochromatin state distinct from both active and silenced chromatin.(28) The chromatin context of Kaiso targets is thus highly heterogeneous, spanning H3K9-methylated, hypoacetylated promoters,(19) H3K4me1-marked compacted chromatin,(28) and acetylated active chromatin at unmethylated targets,(27) suggesting that nucleosome positioning and compaction profoundly influence whether and how Kaiso engages methylated DNA *in vivo*. Yet no study has directly tested whether Kaiso can bind methylated DNA in a nucleosomal context, or how nucleosome packaging affects the affinity and specificity of this interaction. Thus, it remains unclear whether the discrepancy between *in vitro* and *in vivo* binding reflects an intrinsic inability of Kaiso to access nucleosomal DNA, or whether Kaiso can recognize methylated nucleosomes only under conditions that favor partial DNA unwrapping.

Here, we report the first direct evidence that Kaiso binds methylated nucleosome core particles with strong positional preference, engaging full and hemi-methylated CGCG sites at the nucleosome entry/exit region (SHL 6.5) and losing methylation specificity toward the dyad. Solution NMR show that Kaiso binding displaces the H3 N-terminal tail from its native nucleosomal DNA contacts, in a perturbation pattern closely matching the one produced by the pioneer factor Sox2 at a similar region,(29) suggesting that H3 tail displacement is a general consequence of TF binding at the entry/exit sites. These findings provide a mechanistic basis for understanding how Kaiso navigates the chromatin landscape *in vivo* and have implications for how methyl-CpG readers and pioneer factors coordinate DNA methylation readout with histone modification states in transcriptional regulation.

## Methods

### Protein preparation

Kaiso(471–604) WT and E535A constructs containing the optimal C_2_H_2_ zinc finger DNA-binding domain and lacking the BTB/POZ domain were kindly provided by Prof. Peter Wright (The Scripps Research Institute, La Jolla, CA). Kaiso was expressed and purified as previously described.(15) The Kaiso E535A mutant was generated using standard methods for site-directed mutagenesis. The lyophilized protein was dissolved in [10 mM Tris (pH 7.0), 1 mM TCEP] buffer and quantified by the absorbance at 280 nm (NanoDrop, Thermo Scientific). The protein was refolded (∼20 μM final) by gradual addition to the buffer supplemented with 3.3 molar equivalents of Zn^2+^, flash frozen after addition of 10% glycerol, and stored at -80 °C.

### DNA preparation

The palindromic Kaiso binding motif (5′-TCTCGCGAGA-3′) was inserted into the Widom 601 sequence in the pGEM-3z/601 plasmid (a gift from Dr. Greg Bowman, Johns Hopkins University) by standard site-directed mutagenesis. DNA sequences are listed in Table S1. Substrate DNAs (10-60 ml preparative PCR reactions) were generated from this template as previously described,(30), using either unlabeled primers or 5′-fluorophore-labeled primers (6-FAM or Cy3) obtained from IDT with HPLC purification. Primer lengths were chosen to position the Kaiso motif at superhelical locations SHL 6.5, 5.5, 2.5, or 0.5 within the reconstituted nucleosome. Unmethylated substrates were prepared using an unmodified 5′-Cy3 primer, while hemi- and fully methylated substrates were prepared using a 5′-Cy3 primer carrying 5-methylcytosine at the central CGCG of the Kaiso motif (Table S2). PCR products were purified by preparative vertical native gel electrophoresis on a 6% (60:1 acrylamide/bisacrylamide) gel using a 491 Prep Cell (Bio-Rad) run at 11 W and 4 °C as previously described.(31) Eluted fractions were analyzed by agarose gel electrophoresis, pooled, concentrated, and stored at -20 °C. For full methylation, hemi-methylated DNA (265 nM, ∼120 µg) was incubated in a 5 mL reaction containing 200 Units DNMT1 (NEB), 1 mM S-adenosyl-L-methionine (SAM; NEB), 0.1 mg/mL BSA (NEB), and 10 mM DTT in 1X DNMT1 reaction buffer (NEB) at 37 °C for 24 - 48 hrs. Reaction progress was monitored by 5% native TBE-PAGE (125 V, 45 min), as described below. The fully methylated DNA product was purified as described above.

### Validation of substrate methylation status

To assess methylation at the target Kaiso CpGs, 20 µL reactions were prepared by mixing 5 µL of DNMT1-treated DNA with an equal molar amount of FAM-labeled unmethylated DNA in 1X CutSmart buffer (NEB), then either (i) heated to 95 °C for 10 min and slow-cooled at room temperature for 30 min or (ii) left at room temperature without heating. NruI (NEB, 1 µL per reaction) was added, along with one heated reaction containing no enzyme as a control. Reactions were incubated at 37 °C for 1 hr. To assess off-target CpG methylation, 5 µL aliquots of DNA were digested with 0.25 µL of BstUI, AciI, BsaAI, or HpaII (all NEB) in 20 µL reactions in 1X CutSmart buffer at 37 °C for 1 hr. Digestion products (2 µL) for each reaction were resolved on a 5% native TBE-PAGE gel (125 V, 45 min, room temperature) and visualized on a Typhoon 5 imager using FAM or Cy3 fluorescence. Gels were processed using ImageJ2(32) with background subtractions to obtain intensity profiles. Methylation efficiency at the Kaiso motif was calculated from the FAM-channel bands in the NruI cut assay. After heat-and-reanneal of the DNMT1-treated product with unlabeled unmethylated DNA, the FAM-labeled population consists of two equimolar duplex species: the parental DNMT1-treated duplex (A, uncut by NruI) and a hybrid duplex (B) in which the FAM strand pairs with an unmethylated complement, which is cleaved by NruI only when the FAM strand is unmethylated (B_1_). The methylation efficiency *f* on the FAM strand was therefore calculated as *f* = 1 – 2(B_1_/(A + B)), where B_1_ is the integrated intensity of the cut band and (A + B) is the total FAM signal. To obtain the integrated area of the Gaussian, intensity profiles were fit by nonlinear least-squares to a sum of two Gaussians plus a constant baseline, with all six parameters (amplitude, center, and width of each band) allowed to float, using in-house scripts.

### Histone preparation

*Xenopus laevis* (XL) histones H2A, H2B, H3, and H4 in pET3a were a generous gift from Dr. Greg Bowman (JHU). Histone plasmids were transformed into E. coli BL21(DE3) pLysS for expression in 2XTY medium (unlabeled) or M9 minimal medium (for ^15^N isotopic labeling). ^15^N labeling was achieved by supplementing M9 medium with 1 g/L ^15^NH_4_Cl (Cambridge Isotope Laboratories). H2A, H2B, and H3 were purified on a 20 mL tandem Q-SP HiTrap HP column (Cytiva; 2 × 5 mL Q over 2 × 5 mL SP) in sodium acetate-urea buffer as previously described.(30) H4 was recovered from inclusion bodies and purified on a HiPrep 26/10 desalting column (GE Healthcare) followed by a 2xQ-SP HiTrap column (Cytiva) as previously described.(30, 33)

### Nucleosome preparation

Lyophilized histones were resuspended at ∼2 mg/mL in unfolding buffer (20 mM Tris-HCl pH 7.8, 6 M guanidine-HCl, 5 mM DTT) with a ∼20% molar excess of H2A/H2B, then refolded by four sequential dialyses against refolding buffer (10 mM Tris-HCl pH 7.8, 2 M NaCl, 1 mM EDTA, 5 mM β-mercaptoethanol (BME)) at 4°C using 3.5 kDa MWCO membranes (Spectra Labs). Refolded octamer and excess H2A-H2B dimer were separated on a HiLoad 16/600 Superdex 200 pg column (Cytiva) as previously described.(31, 33) Fractions were analyzed by SDS-PAGE, concentrated to 50-150 µM, flash-frozen, and stored at -80 °C. Nucleosomes were assembled by an adapted salt-dialysis protocol,(31, 33) combining octamer (6 µM), H2A-H2B dimer (1.8 µM), and 10% molar excess of DNA (6.6 µM). Reconstituted nucleosomes were purified by preparative vertical native gel electrophoresis on a 7% (60:1 acrylamide/bisacrylamide) gel, as previously described.(30) Nucleosome purity was assessed by 5% native TBE-PAGE.

### Electrophoretic mobility shift assay (EMSA)

5′-FAM/5′-Cy3-labeled nucleosome (10 nM, NCP) was mixed with variable concentrations of Kaiso WT or E535A (3 nM to 1000 nM) in binding buffer (10 mM Tris-HCl pH 7.5, 150 mM KCl, 1 mM DTT, 0.5 mM TCEP, 1 mg/ml Non-Fat Dry Milk (Bio-Rad), 0.05% Tween-20, 8% glycerol) in a total reaction volume of 20 µL and incubated at 25 °C for 30 min. Reactions were loaded (2 µL) on a 5% native polyacrylamide gel and run for 90 min at 125 V on ice. EMSAs were acquired with 3-5 replicates each. Binding was visualized using a Typhoon 5 bioimager (GE Amersham) using FAM fluorescence. Gels were processed using ImageJ2(32) with background subtractions. The integrated peak intensities for the specific complex (I_spec_), non-specific complex (I_ns_), and the free nucleosome (I_free_) were quantified for each lane. Because quantification at 0 nM Kaiso yielded non-zero signal in the specific and non-specific regions (from free-band tails, background, etc.), we performed a per-gel, per-protein background subtraction at the raw-intensity level: for each gel and protein, the intensities I_spec_(0) and I_ns_(0) in the absence of Kaiso were subtracted from the corresponding regions of every other lane in that set. The corrected Kaiso (=P) fractional binding was then *f*_bound, total_(P) = [I_spec, corr_(P) + I_ns, corr_(P)] / [I_free_(P) + I_spec, corr_(P) + I_ns, corr_(P)]. Each titration was fit to a quadratic 1:1 binding model with ligand depletion, *f*_bound, total_(P) = *f*_max_ x [(K_d_ + P + N − ((K_d_ + P + N)^2^ − 4P x N))^1/2^ / (2N)], with N = 10 nM fixed and (K_d_, *f*_max_) as free parameters using an in-house python script. Per-gel outliers were identified using Tukey’s 1.5xIQR rule, a 3-fold-from-median ratio rule, and rejection of fits with SE(K_d_) > K_d_; outliers were removed only if n ≥ 3 replicates remained. K_d_ values are reported as mean ± SD across non-outlier replicates. In addition, to directly assess the population of the discrete shifted band, we extracted the specific complex fraction (*f*_spec_(100 nM) = I_spec_ / (I_free_ + I_spec_ + I_ns_), using background- and loading-corrected intensities) within a fixed band-position window defined on a methylated lane and applied identically to all lanes on the same gel. Specific complex fraction analysis was based on the same gels selected for K_d_ analysis.

### NMR spectroscopy

An ^15^N-H3 MeCG_2_hemi (SHL6.5) nucleosome sample was prepared and dialyzed 3 times against NMR buffer (20 mM Tris-HCl pH 7.0, 25 mM NaCl, 0.5 mM TCEP) using a 3.5 kDa MWCO membrane (Spectra Labs), then concentrated to ∼20 µM (∼300 µL) in an Amicon Ultra-0.5 (10 kDa MWCO) centrifugal concentrator at 4 °C. For Kaiso titrations, 50 µM unlabeled Kaiso DBD was added slowly to the nucleosome at the desired molar ratios, and the sample was reconcentrated to ∼300 µL after each addition. NMR buffer was supplemented with 7.5% D_2_O (Cambridge Isotope Laboratories) prior to data collection. Spectra were acquired on a Bruker Ascend 800 MHz spectrometer equipped with a triple-resonance cryoprobe. Standard 2D ^1^H-^15^N HSQC experiments (64 scans) were collected at 37 °C. Data was processed using nmrPipe(34) and analyzed using Sparky (Goddard, T. D. & Kneller, D. G. SPARKY 3, University of California, San Francisco) and in-house scripts. Chemical shift perturbations (CSPs) were calculated from the difference in proton (Δδ_H_) and nitrogen (Δδ_N_) chemical shifts in HSQC spectra, as follows: CSP = (Δδ_H^2^_+ (0.154Δδ_N_)^2^)^1/2^.

## Results

### Preparation of site-specifically CpG methylated nucleosomes

Kaiso has been shown to associate with primed heterochromatin in a DNA-methylation-dependent manner.(28) To assess the binding of Kaiso to CpG-methylated nucleosomes, we prepared constructs with a Widom 601 sequence modified with the *in vivo* consensus Kaiso binding sequence(35) (MeCG_2_) at different positions (Figure 1A, Table S1). Crystallographic studies have shown that the Kaiso DBD engages this site as a monomer through three tandem zinc fingers (ZF1-ZF3, Figure 1C).(17) To ensure that this binding interface remains accessible in the nucleosomal context, we positioned the MeCG_2_ site at SHL 0.5, 2.5, 5.5, or 6.5, placing the predicted major groove of the Kaiso recognition site outward-facing relative to the histone octamer (Figure 1B, Figure 1D). For each construct, we prepared DNA that was unmethylated or CpG methylated on one strand (hemi-methylated) of the Kaiso motif. For SHL 5.5 and 6.5 constructs, we also prepared fully methylated DNA on both strands of the Kaiso motif. We synthesized the site-specifically hemi-methylated 601 DNA by using polymerase chain reaction (PCR) with one primer containing methylated cytosines at the Kaiso motif (Figure 1B). To make the fully methylated DNA, we then modified enzymatically hemi-methylated DNA using the methyltransferase DNMT1, which methylates cytosines selectively in hemi-methylated CpG steps (Figure 1B).(36)

**Figure 1.**
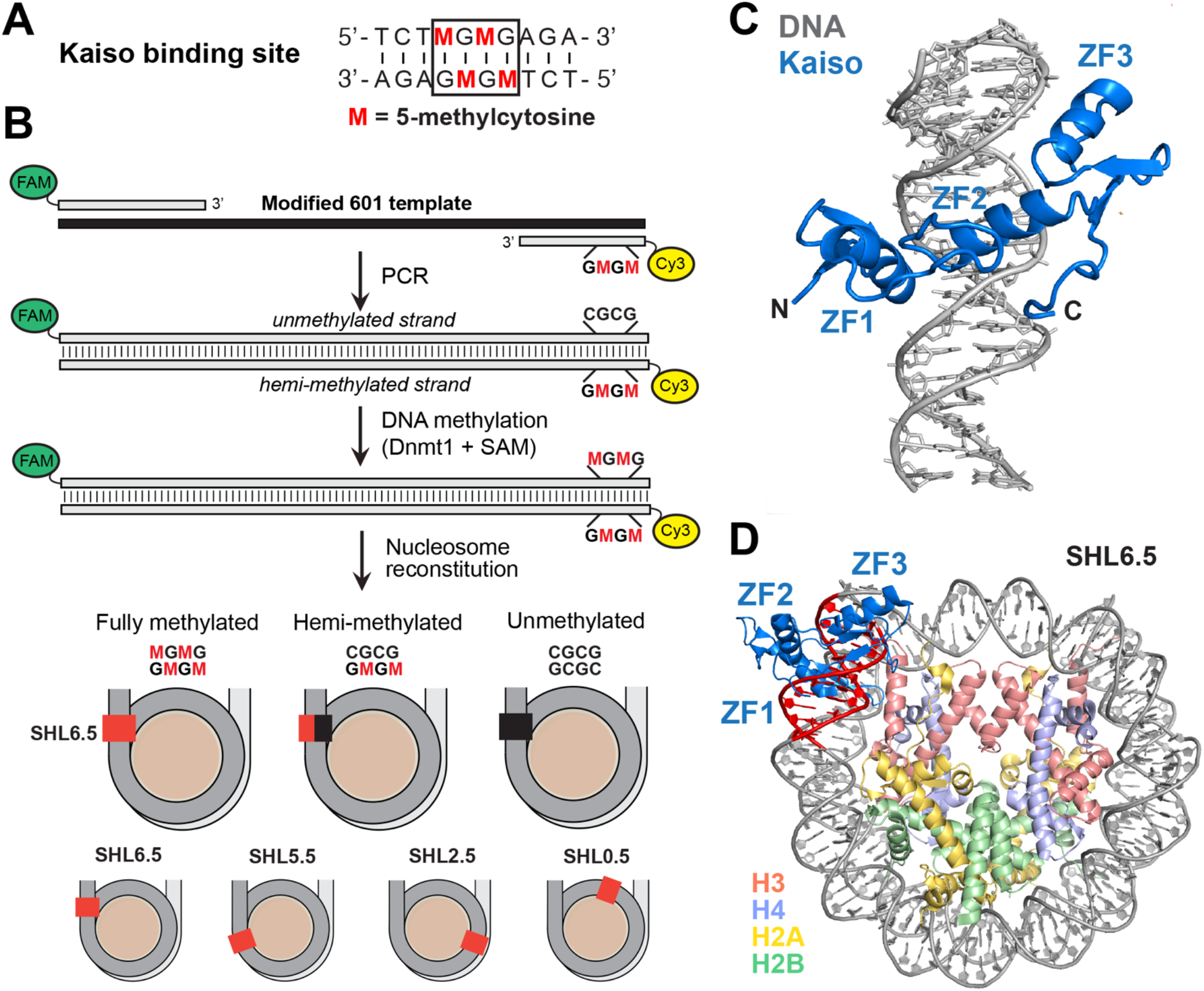
Design of methylated nucleosome substrates for Kaiso binding studies. **(A)** The Kaiso recognition motif (MeCG_2_) with the central palindromic CGCG core (boxed). The two CpG dinucleotides at the core can be methylated to 5-methylcytosine (M, red), defining fully methylated, hemi-methylated (one strand), and unmethylated states. **(B)** Substrate preparation strategy. The Kaiso binding site, present in a modified Widom 601 plasmid template, was amplified by PCR with 5′ FAM- and Cy3-labeled primers to generate a 147 bp duplex with the two fluorophores at opposite ends. Pre-methylated cytosines on the Cy3-labeled primer yield the hemi-methylated substrate. Subsequent DNMT1/SAM treatment converts it to the fully methylated state, while PCR with unmodified primers gives the unmethylated control. Substrates were reconstituted into nucleosome core particles (NCPs) with the Kaiso binding site positioned at SHL 6.5 in three methylation states (top row), or with the site alternatively placed at SHL 5.5, 2.5, or 0.5 (bottom row). Red squares mark the approximate position of the Kaiso binding site within each nucleosome. **(C)** Crystal structure of the Kaiso DBD (blue) bound to the MeCG_2_ recognition sequence shown in (A) (PDB 5VMV), with the three zinc fingers labeled ZF1, ZF2, and ZF3. **(D)** Model of Kaiso (blue) bound to the methylated CGCG site in (A) (red) placed at SHL 6.5 within a modified Widom 601 nucleosome (PDB 3LZ0).

To assess the extent of DNA methylation, we used the restriction enzyme NruI, which makes a blunt cut at the center of TCG|CGA motifs and whose activity is blocked by cytosine methylation (Figure 2A,C). Since NruI does not cut hemi-methylated DNA, we measured the extent of full methylation by mixing DNMT1 products with corresponding unmethylated DNA in 1:1 ratio, heating to 95 °C, and cooling to allow for the single strands to randomly reshuffle upon annealing (Figure 2A). We set up reactions, where the DNMT1 product DNA was 5’-FAM and 5’-Cy3 labeled on each side, with the fully unmethylated DNA unlabeled. If the DNA was 100% methylated by DNMT1, we anticipated that 100% of the total FAM-labeled DNA would contain a methylated strand after reshuffling and be resistant to NruI cutting. If the DNA was 0% methylated by DNMT1, then 50% of FAM-labeled DNA would be resistant to NruI cleavage. Using this method, we estimated that the fully methylated DNA products in large scale reactions varied from 64% (SHL 6.5 DNA) to 83% (SHL 5.5 DNA) (Figure 2C). As controls, we set up reactions where no NruI was added but the strands were reshuffled (lane 1) and where the strands were not reshuffled but NruI was added (lane 3) (Figure 2C).

**Figure 2.**
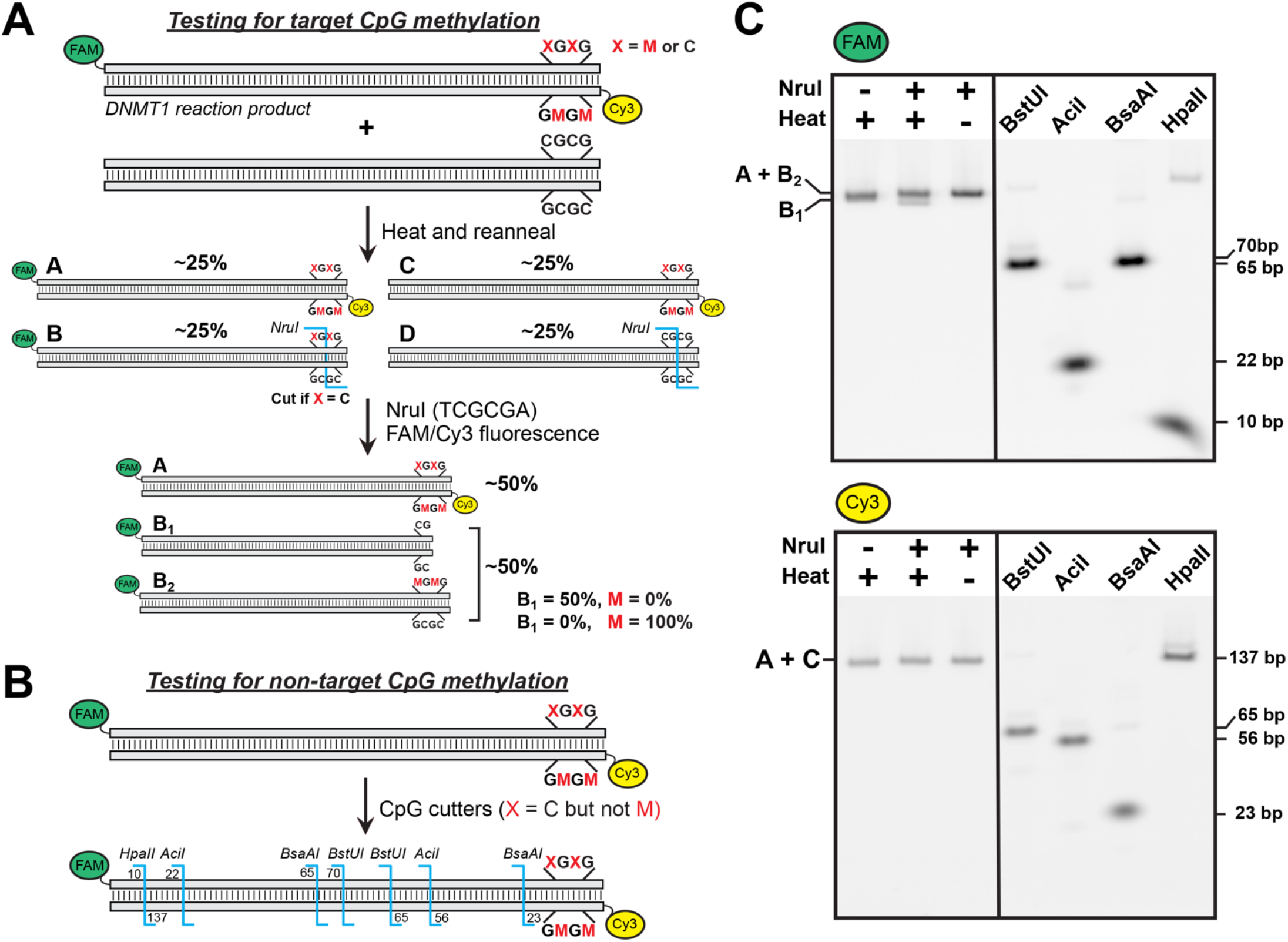
Validation of on-target methylation and off-target specificity of the MeCG_2_ SHL 6.5 substrate. **(A)** Schematic of the NruI cut assay for target-CpG methylation. The DNMT1 reaction product is mixed 1:1 with unmethylated unlabeled DNA and reannealed, producing four equimolar duplex species (A–D). Subsequent NruI digestion cuts only molecules, in which the FAM-strand CpG is unmethylated. The fraction of FAM signal in the cut “B_1_” band reports directly on DNMT1 methylation efficiency, ranging from 50% (no DNMT1 activity) to 0% (complete methylation). **(B)** Strategy for testing non-target CpG methylation. Predicted FAM- and Cy3-labeled fragment sizes are shown for digestion with four methylation-sensitive enzymes (HpaII, BsaAI, AciI, BstUI) at their respective sites within the MeCG_2_ substrate. **(C)** Native PAGE gels for the FAM (top) and Cy3 (bottom) channels. Left: NruI assay following heat-and-reanneal. Right: digestion with non-target CpG cutters at predicted sizes, confirming that DNMT1-mediated methylation is restricted to the Kaiso-site CpG. M = 5-methylcytosine; X = M or C.

We also ensured that DNMT1 did not methylate other CpG sites in different sequence contexts in the modified 601 DNA. This was done by subjecting the product of the DNMT1 reaction to four other restriction enzymes, which cut DNA only at unmethylated CpG steps with different sequence specificity (Figure 2B). The enzymes were BstUI (5’-CG|CG-3’), AciI (5’-C|CGC-3’), BsaAI (5’-YAC|GTR-3’), and HpaII (5’-C|CGG-3’), each with one or more target sites within the 601 sequence (Figure 2B). In each case, nearly complete digestion (i.e. the shortest product resulting from multiple cut sites) of the DNMT1 product was obtained with all four enzymes, suggesting there was no observable off-target methylation by DNMT1. By contrast, the same digestion test of DNA modified by the methyltransferase M.SssI, which indiscriminately methylates cytosines at CpG steps, was severely inhibited (data not shown). Thus, using this method we were able to prepare site-selectively CpG-methylated 147 base-pair long DNA to be used for nucleosome preparation.

### Kaiso binds methylated nucleosomes preferentially at the entry/exit site

To dissect how Kaiso recognizes methylated CpGs in a nucleosomal context, we performed electrophoretic mobility shift assays (EMSAs) under near-physiological salt conditions on nucleosomes containing the Kaiso binding motif at SHL 0.5, 2.5, 5.5, or 6.5, with the central CpG steps either unmethylated, hemi-methylated, or fully methylated. Across the full Kaiso titration (0-100 NCP:Kaiso ratios), unmethylated nucleosomes at all positions produced only a diffuse, smeared supershift, characteristic of heterogeneous low-affinity engagement. In contrast, hemi- and fully methylated substrates at SHL 6.5 yielded a stable, discrete shifted complex migrating just above the free nucleosome that grew with increasing Kaiso concentrations and saturated within the titration range. Methylation-dependent binding was most pronounced at SHL 6.5. Moving the Kaiso site 10 base pairs inward to SHL 5.5 still produced a methylation-dependent shift but with a smaller dynamic range. At dyad-proximal positions (SHL 0.5 and 2.5), no methylation-dependent enhancement was observed, indicating that Kaiso cannot productively engage the recognition motif when it is buried against the histone octamer. As a control for methyl-CpG-specific binding, we tested the E535A mutant, which we previously showed disrupts canonical and non-canonical C–H⋯O hydrogen bonds with the 5-methylcytosine (5mC) nucleobase and decreases binding affinity in the context of free DNA.(17) Kaiso E535A failed to form the discrete shifted complex on methylated substrates, instead producing a smeared supershift resembling that of Kaiso WT on unmethylated DNA, consistent with loss of specific methylation readout while retaining residual non-specific DNA binding.

**Table 1.** Binding affinities (K_d_) for Kaiso WT and E535A with nucleosome substrates derived from EMSA.

| SHL | NCP Substrate | Kaiso WT<br>$K_d(\text{total}) \pm \text{SD} (n^*)$ (nM) | Kaiso E535A<br>$K_d(\text{total}) \pm \text{SD} (n)$ (nM) |
| --- | --- | --- | --- |
| 6.5 | MeCG<sub>2</sub>full | 98 ± 15 (3) | 340 ± 102 (3) |
| 6.5 | MeCG<sub>2</sub>hemi | 104 ± 22 (4) | 243 ± 122 (5) |
| 6.5 | CG<sub>2</sub> | 370 ± 158 (3) | 309 ± 136 (3) |
| 5.5 | MeCG<sub>2</sub>full | 169 ± 26 (5) | 138 ± 56 (3) |
| 5.5 | MeCG<sub>2</sub>hemi | 318 ± 122 (4) | 254 ± 126 (5) |
| 5.5 | CG<sub>2</sub> | 513 ± 137 (3) | 192 ± 22 (3) |

\*n = number of independent measurements

Quantitative fitting of the total fraction bound revealed that at SHL 6.5, Kaiso WT bound fully methylated (K_d_ = 98 ± 15 nM) and hemi-methylated (K_d_ = 104 ± 22 nM) substrates with comparable affinity, roughly 4-fold tighter than unmethylated (K_d_ = 370 ± 158 nM). We note that the fully methylated SHL 6.5 substrate was only ∼64% complete in DNMT1-mediated methylation, so the apparent affinity difference between hemi- and fully methylated nucleosomes likely underestimates the true difference. The comparable K_d_ values nevertheless indicate that a single methylated CpG on the strand accessible to E535 is sufficient for high-affinity engagement, with the methyl group on the complementary strand contributing only modestly. This is consistent with the affinity trend on free DNA from our prior bio-layer interferometry (BLI) measurements, where hemi-methylated substrates bound Kaiso only ∼3-fold weaker than fully methylated and ∼60-fold tighter than unmethylated, with specificity dominated by E535 interaction with a single 5mC on one strand.(17) At SHL 5.5, the methylation-dependent discrimination was preserved but reduced in magnitude (K_d_ = 169 ± 26, 318 ± 122, and 513 ± 137 nM for fully, hemi-, and unmethylated, respectively), indicating attenuated recognition at the more internal position. The E535A mutation, which abolishes canonical and C–H⋯O hydrogen bonds and reduces methylated-CpG affinity to near-unmethylated levels on free DNA,(17) produced a position- and methylation-dependent loss of affinity. Specifically, a ∼3.5-fold penalty on fully methylated SHL 6.5 (K_d_ = 340 ± 102 nM), a ∼2.3-fold penalty on hemi-methylated SHL 6.5 (K_d_ = 243 ± 122 nM), and essentially no effect on unmethylated SHL 6.5 or SHL 5.5 substrates was observed. This pattern confirms that the same E535–5mC interaction underlying free DNA recognition is operative on the nucleosome, with its energetic contribution most fully realized at the accessible entry/exit region.

To complement the total apparent affinity analysis, we directly quantified the population of the discrete shifted complex at 100 nM Kaiso (10:1 Kaiso:NCP), where this band reaches its maximum on the gel. At SHL 6.5, WT Kaiso captured ∼42–44% of nucleosomes in the discrete complex on hemi- and fully methylated substrates, nearly 3-fold the unmethylated baseline (∼16%); E535A reduced this to 13–21% across all methylation states, abolishing methylation-dependent specificity. At SHL 5.5, only fully methylated nucleosomes produced a clear enhancement of the specific complex (30% versus 14% on unmethylated for WT), with hemi-methylated nucleosomes indistinguishable from unmethylated.

Together with the affinity measurements, these data show that Kaiso binds methylated nucleosomal DNA with high affinity and specificity primarily at the entry/exit region, that hemi-methylation is sufficient for full engagement when the recognition motif is positioned at SHL 6.5 in the optimal orientation, and that E535 is required for methyl-CpG-specific recognition in this context.

### Kaiso binding at the nucleosome edge displaces the histone H3 N-terminal tail

Because Kaiso binds near the nucleosome entry/exit site at SHL 6.5, we asked whether this interaction perturbs the H3 N-terminal tail. Unlabeled Kaiso was titrated (0, 0.5, 1, 2, and 4X molar excess) into 20 µM ^15^N-labeled H3 nucleosomes hemi-methylated at SHL 6.5, and chemical shift perturbations (CSPs) were measured by solution NMR for the visible H3 tail residues (Figure 4). Kaiso binding produced a discontinuous perturbation pattern, with significant CSPs clustered at residues T3-R8 and A15-A29 and intervening residues remaining near baseline. A subset of peaks split into two populations upon Kaiso addition (Figure 4), consistent with a single Kaiso molecule binding one face of the nucleosome and selectively perturbing the H3 tail on that face while leaving the tail on the opposite face unperturbed. The direction of the chemical shift changes typically matched those reported previously for the free H3 peptide,(37, 38) indicating that the perturbed tail is released from its nucleosomal DNA contacts rather than redirected into a new ordered interaction with Kaiso.

**Figure 3.**
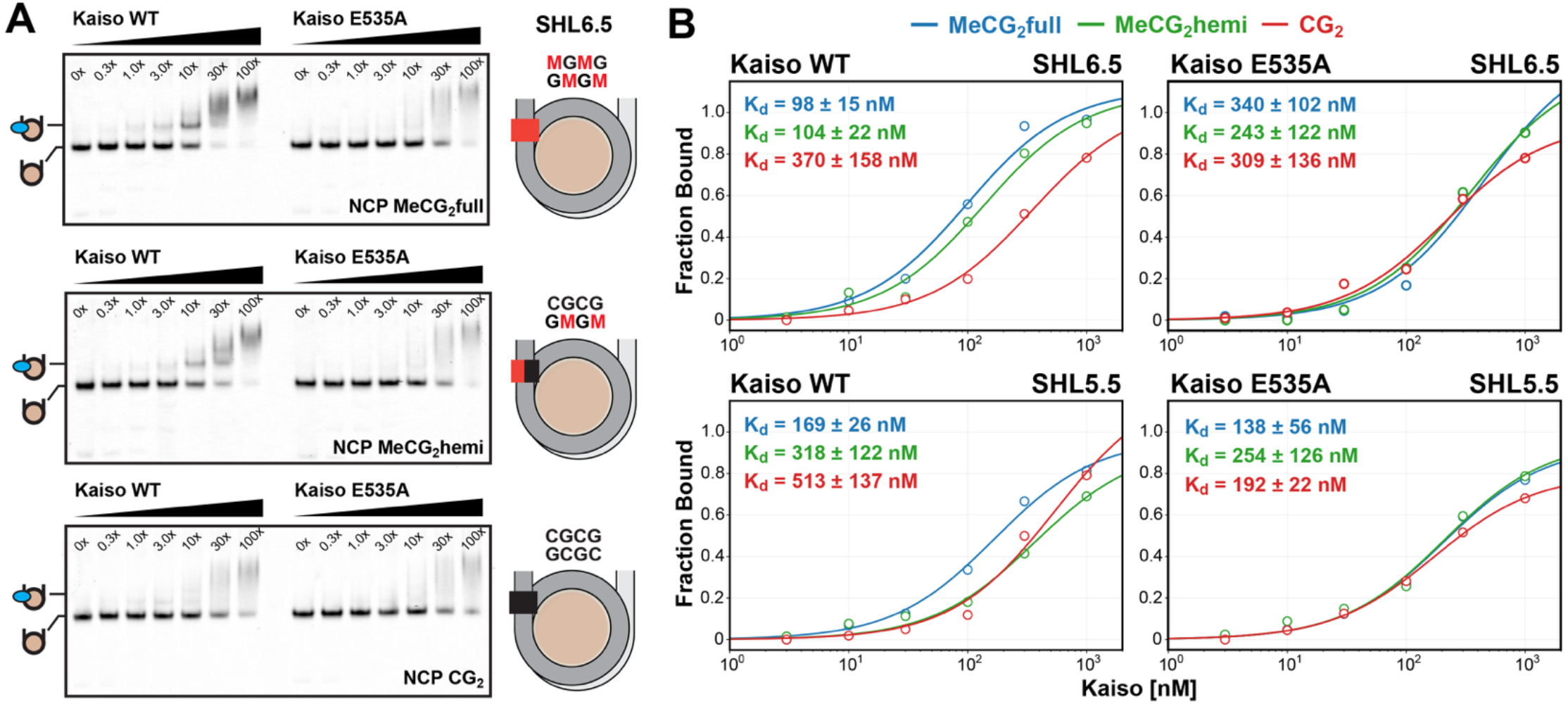
Kaiso binding to methylated nucleosomes is position- and methylation-dependent. **(A)** Representative EMSA gels of WT and E535A Kaiso (0, 0.3, 1, 3, 10, 30, and 100x) titrated against 10 nM fully methylated (NCP MeCG_2_full, top), hemi-methylated (NCP MeCG_2_hemi, middle), and unmethylated (NCP CG_2_, bottom) nucleosomes with the Kaiso binding site at SHL 6.5. Schematics on the right indicate the methylation state of the central CpG steps at the Kaiso site in each substrate. **(B)** Representative apparent binding curves derived from quantification of the EMSA gels for WT (left) and E535A (right) Kaiso at SHL 6.5 (top) and SHL 5.5 (bottom). Fraction bound was calculated for each Kaiso concentration and fit to a 1:1 quadratic binding model accounting for ligand depletion. Mean K_d_ values ± SD across 3-5 independent gel replicates are indicated on each panel. M = 5-methylcytosine.

**Figure 4.**
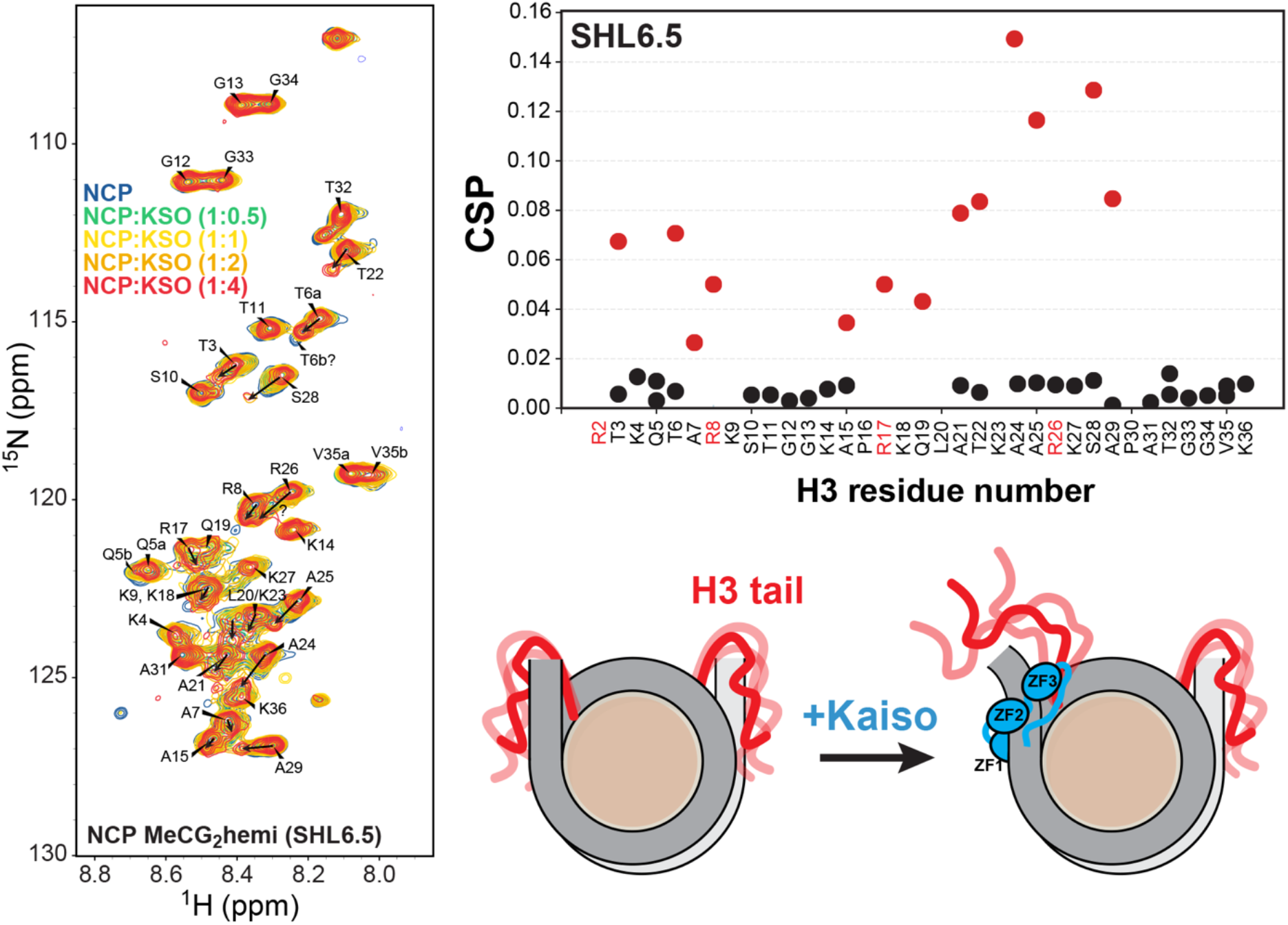
Kaiso binding perturbs the H3 N-terminal tail at the nucleosome entry/exit site. **(A)** ^1^H-^15^N HSQC spectra of ^15^N-H3 NCP MeCG_2_hemi SHL 6.5 titrated with Kaiso (0.5, 1, 2, and 4X). **(B)** Per-residue chemical shift perturbations (CSPs) at 1:4 NCP:Kaiso, showing the appearance of a second population (red circles) consistent with H3 tail displacement near the Kaiso binding site. Arg residues known to make persistent DNA contacts (R2, R8, R17, R26) are highlighted in red on the x-axis. R26 CSP could not be reliably determined. **(C)** Model for Kaiso engagement at SHL 6.5: binding of the Kaiso zinc finger DBD near the entry/exit site displaces the H3 N-terminal tail from its native nucleosomal DNA contacts and likely deforms the DNA helix away from the histone octamer, based on the published Kaiso-MeCG_2_ complex (PDB 5VMV).

The perturbation hotspots cluster around H3 tail Arginine residues that prior NMR studies(37-39) and our published molecular dynamics (MD) simulations independently identified as making persistent DNA contacts near the entry/exit site (R2, R8, R17, R26).(29) This correspondence supports a model, in which Kaiso binding releases the tail specifically from the DNA segment it occupies, rather than inducing a global rearrangement. The Kaiso CSP fingerprint also closely resembles the pattern we reported for the pioneer factor Sox2 bound near SHL 6, which perturbs an overlapping set of residues to a similar extent (Figure S2).(29) Thus, we observe that two structurally unrelated DNA binders, a methylation-dependent major-groove binding zinc-finger repressor and an HMG-box minor-groove binding pioneer factor, produce similar perturbation patterns at this nucleosomal region. This convergence suggests that H3 tail displacement is a general consequence of factor binding at the entry/exit site, driven by steric competition for the same DNA segment rather than by protein-specific contacts.

## Discussion

Our results show that Kaiso binds methylated CGCG sites with high affinity when they are positioned at the nucleosome entry/exit region (SHL 6.5), with reduced affinity at the more internal positions (SHL 5.5), and with no methylation-dependent enhancement at dyad-proximal positions (SHL 0.5 and 2.5). At SHL 6.5, hemi- and fully methylated substrates bind Kaiso with essentially the same affinity, indicating that methylation on one strand is sufficient for high-affinity recognition. This positional behavior identifies Kaiso as a sequence-specific DNA-binding protein, similar to some pioneer TFs,(40, 41) that depends on transient exposure of its target sites through nucleosomal breathing for productive engagement.(30, 42-45)

The SHL-dependent affinity hierarchy we observe (full ≈ hemi > unmethylated) reflects a fundamental access constraint on methyl-CpG readout in the chromatin context. Kaiso’s three zinc fingers must engage ∼10 base pairs of contiguous DNA,(15, 17) a footprint not continuously exposed on the wrapped nucleosome. Productive binding therefore requires transient DNA unwrapping at the entry/exit region. Kaiso’s intrinsic tendency to bend bound DNA toward ZF2(17) would further aid this process by deflecting the nucleosomal DNA away from the histone surface, promoting local DNA unwrapping. The E535A mutation produces its largest penalty on methylated substrates at SHL 6.5 and little effect at internal positions, where DNA accessibility itself limits engagement. This pattern confirms that the same E535–mCpG hydrogen bonds driving free-DNA recognition(17) also operate within the chromatin context. The near-equivalent affinity of hemi- and fully methylated substrates at SHL 6.5 extends a striking feature of Kaiso’s free-DNA recognition mechanism into chromatin. Our previous BLI measurements showed that hemi-methylated free DNA binds Kaiso only ∼3-fold weaker than fully methylated, while unmethylated DNA is ∼60-fold weaker, with specificity dominated by a single 5^′^ mCpG contacted by E535.(17) Although Kaiso’s recognition motif TCTCGCGAGA is palindromic at the central CGCG, our crystal structures showed that Kaiso engages it in a single defined orientation set by the asymmetric flanking sequence, with a 5^′^ C-G favoring the productive orientation.(17, 46) Our nucleosomal substrates were designed to align this preferred orientation with the geometry at SHL 6.5, placing the dominant 5^′^ mCpG in the outward-facing major groove. The near-equivalent hemi/full affinity at this position therefore indicates that the second methyl contributes little additional binding energy on the nucleosome. This model predicts that reversing the methylation strand in a hemi-methylated substrate would reduce Kaiso affinity and could shift its binding orientation.

The biological significance of Kaiso’s hemi-methylation tolerance extends beyond the biophysics of methyl-CpG readout. Hemi-methylated CpG is the intermediate state that exists immediately behind the replication fork, when the parental strand carries the methylcytosine mark but the newly synthesized daughter strand does not, and this state persists until DNMT1 completes maintenance methylation.(47, 48) Kaiso’s ability to bind hemi-methylated nucleosomes therefore raises the possibility that it could function as a replication-coupled reader, preserving transcriptional repression at methylated loci through S phase before maintenance methylation is complete.(49) By analogy with UHRF1, which recognizes hemi-methylation to recruit DNMT1,(50-52) Kaiso could provide an independent mechanism for heritable silencing that buffers against the natural delay in DNMT1 activity.(53) To our knowledge, this role has not been directly tested for Kaiso. Direct demonstration would require showing that Kaiso travels with the replisome and remains bound to nascent hemi-methylated chromatin in cells.

Beyond the affinity measurements, the most mechanistically striking result of this work is the close match between our Kaiso NMR perturbation fingerprint and the one we recently reported for the pioneer factor Sox2.(29) Despite recognizing DNA through fundamentally different chemistries, where Sox2 HMG DNA-binding binds and widens the DNA minor groove,(54, 55) the two proteins produce nearly identical patterns of H3 tail perturbation when bound near the nucleosome edge. This convergence suggests that H3 tail displacement is not a property of any protein or recognition mode, but a general consequence of factor binding at entry/exit DNA and competing with the H3 N-terminal tail for the same DNA interface. The displacement model is supported by two complementary observations: the residues most perturbed by Kaiso binding correspond to those that make persistent DNA contacts in previously published MD simulations and NMR analyses of the H3 tail in the apo nucleosome,(29, 37, 39, 56, 57) and the perturbed peaks shift toward chemical shift values characteristic of the free H3 tail peptide,(37, 38) consistent with release of the tail from its nucleosomal DNA contacts. Direct structural tests will require MD simulations with Kaiso explicitly bound at SHL 6.5, potentially complemented by cross-linking mass spectrometry to map Kaiso-H3 tail proximities.

The consequences of H3 tail displacement likely extend beyond the immediate binding event. The H3 tail is the substrate for the principal histone-modifying enzymes,(58, 59) and contributes to the inter-nucleosomal contacts that drive chromatin compaction.(60-62) Tail displacement induced by factor binding could therefore function as a local “unmasking” event,(37, 63, 64) exposing the tail for covalent modification and loosening compaction enough for access by additional factors. For Kaiso specifically, this reframes its role from a passive methyl-CpG sensor into an active local destabilizer of methylated chromatin. Genome-wide analyses have associated methylated Kaiso binding sites with a distinctive chromatin signature: compacted regions carrying H3K4 monomethylation in the absence of other active or repressive marks, a “primed heterochromatin” state distinct from canonical heterochromatin.(28) The same study did not detect the long-proposed Kaiso-NCoR-HDAC complex(19) genome-wide at these sites, suggesting Kaiso’s chromatin activity may be more intrinsic than co-repressor-mediated.(28) Sox2 binding sites are similarly enriched for H3K4me1 *in vivo* at poised enhancers,(65-67) and the shared association of Kaiso and Sox2 with H3K4me1 chromatin parallels the tail-displacement mechanism we observe biochemically. Tail displacement may therefore be a common step licensing H3K4 monomethylation, with the surrounding compaction limiting further modification. Although direct causality has not been established, our findings suggest Kaiso may make the tail accessible to MLL3/4 monomethyltransferase activity(68-70) by analogy with established cases of tail displacement enhancing modifiability.(37, 71) Testing this will require biochemical assays with the relevant enzymes and cellular experiments asking whether Kaiso is required for H3K4me1 at its methylated targets.

Taken together, these findings provide a mechanistic picture of how Kaiso might operate in its native chromatin context. Kaiso’s major genomic targets are methylated CpG islands, imprinted control regions, and other compacted, nucleosome-dense regions where access by most TFs is restricted.(20, 28, 72) The positional sensitivity, hemi-methylation tolerance, and tail-displacement activity we characterize here collectively explain how Kaiso engages such regions. Entry/exit-biased binding lets Kaiso reach its target CpGs through spontaneous nucleosomal breathing,(42, 43, 73) without requiring prior chromatin remodeling. Tolerance for hemi-methylation supports continuous occupancy across the cell cycle, including the period after DNA replication.(53) And H3 tail displacement provides a local structural amplifier that turns a single recognition event into broader consequences for the surrounding chromatin.(63, 71) This model presents Kaiso as an actively engaged regulator that locally modulates the structure of methylated loci rather than a static heterochromatin resident, consistent with the “primed” chromatin state observed *in vivo* at its methylated targets.(28) The mononucleosome context we use here does not capture the higher-order chromatin environment in which Kaiso ultimately operates, and extending this analysis to reconstituted chromatin arrays or in-cell NMR will be essential to test whether the mechanisms we propose scale from a single nucleosome to a silenced domain.

## Acknowledgements

We thank Prof. Greg Bowman (JHU) for kindly providing materials and access to instrumentation. We acknowledge Helen Moos for assistance with sample preparation. We are grateful to Dr. Ananya Majumdar, director of the Biomolecular NMR center (JHU), for assistance with NMR data collection and to the Integrated Imaging Center (JHU) staff for bioimager training and support. This research was supported by NIGMS grant R01GM147642 awarded to E.N.N.

## Author Contributions

E.N.N. conceptualized the study, designed the experiments, performed sample preparation, EMSA, NMR experiments, data analysis, prepared figures and wrote the manuscript. F.C.M.G. contributed to sample preparation, EMSA, and data analysis.

## Supplementary Data

Supplementary Data are available online.

## References

1. Smith ZD, Meissner A. DNA methylation: roles in mammalian development. Nat Rev Genet. 2013;14(3):204–20.

2. Breiling A, Lyko F. Epigenetic regulatory functions of DNA modifications: 5-methylcytosine and beyond. Epigenetics & Chromatin. 2015;8.

3. Greenberg MVC, Bourc’his D. The diverse roles of DNA methylation in mammalian development and disease. Nat Rev Mol Cell Biol. 2019;20(10):590–607.

4. Bannister AJ, Kouzarides T. Regulation of chromatin by histone modifications. Cell Res. 2011;21(3):381–95.

5. Millan-Zambrano G, Burton A, Bannister AJ, Schneider R. Histone post-translational modifications - cause and consequence of genome function. Nat Rev Genet. 2022;23(9):563–80.

6. Liu R, Wu J, Guo H, Yao W, Li S, Lu Y, et al. Post-translational modifications of histones: Mechanisms, biological functions, and therapeutic targets. MedComm (2020). 2023;4(3):e292.

7. Deng S, Feng Y, Pauklin S. 3D chromatin architecture and transcription regulation in cancer. J Hematol Oncol. 2022;15(1):49.

8. Portillo-Ledesma S, Chung S, Hoffman J, Schlick T. Regulation of chromatin architecture by transcription factor binding. Elife. 2024;12.

9. Yin Y, Morgunova E, Jolma A, Kaasinen E, Sahu B, Khund-Sayeed S, et al. Impact of cytosine methylation on DNA binding specificities of human transcription factors. Science. 2017;356(6337).

10. Reiter F, Wienerroither S, Stark A. Combinatorial function of transcription factors and cofactors. Curr Opin Genet Dev. 2017;43:73–81.

11. Klemm SL, Shipony Z, Greenleaf WJ. Chromatin accessibility and the regulatory epigenome. Nat Rev Genet. 2019;20(4):207–20.

12. Zaret KS. Pioneer Transcription Factors Initiating Gene Network Changes. Annu Rev Genet. 2020;54:367–85.

13. Moyung K, Li Y, MacAlpine HK, Hartemink AJ, MacAlpine DM. Genome-wide nucleosome and transcription factor responses to genetic perturbations reveal chromatin-mediated mechanisms of transcriptional regulation. Genome Res. 2026;36(1):115–28.

14. Daniel JM, Reynolds AB. The catenin p120(ctn) interacts with Kaiso, a novel BTB/POZ domain zinc finger transcription factor. Molecular and cellular biology. 1999;19(5):3614–23.

15. Buck-Koehntop BA, Stanfield RL, Ekiert DC, Martinez-Yamout MA, Dyson HJ, Wilson IA, et al. Molecular basis for recognition of methylated and specific DNA sequences by the zinc finger protein Kaiso. Proceedings of the National Academy of Sciences of the United States of America. 2012;109(38):15229–34.

16. Filion GJ, Zhenilo S, Salozhin S, Yamada D, Prokhortchouk E, Defossez PA. A family of human zinc finger proteins that bind methylated DNA and repress transcription. Molecular and cellular biology. 2006;26(1):169–81.

17. Nikolova EN, Stanfield RL, Dyson HJ, Wright PE. CH…O Hydrogen Bonds Mediate Highly Specific Recognition of Methylated CpG Sites by the Zinc Finger Protein Kaiso. Biochemistry. 2018;57(14):2109–20.

18. Nikolova EN, Stanfield RL, Dyson HJ, Wright PE. A Conformational Switch in the Zinc Finger Protein Kaiso Mediates Differential Readout of Specific and Methylated DNA Sequences. Biochemistry. 2020;59(20):1909–26.

19. Yoon HG, Chan DW, Reynolds AB, Qin J, Wong J. N-CoR mediates DNA methylation-dependent repression through a methyl CpG binding protein Kaiso. Molecular cell. 2003;12(3):723–34.

20. Lopes EC, Valls E, Figueroa ME, Mazur A, Meng FG, Chiosis G, et al. Kaiso contributes to DNA methylation-dependent silencing of tumor suppressor genes in colon cancer cell lines. Cancer research. 2008;68(18):7258–63.

21. Kelly KF, Spring CM, Otchere AA, Daniel JM. NLS-dependent nuclear localization of p120ctn is necessary to relieve Kaiso-mediated transcriptional repression. J Cell Sci. 2004;117(Pt 13):2675–86.

22. Park JI, Kim SW, Lyons JP, Ji H, Nguyen TT, Cho K, et al. Kaiso/p120-catenin and TCF/beta-catenin complexes coordinately regulate canonical Wnt gene targets. Developmental cell. 2005;8(6):843–54.

23. Kim SW, Park JI, Spring CM, Sater AK, Ji H, Otchere AA, et al. Non-canonical Wnt signals are modulated by the Kaiso transcriptional repressor and p120-catenin. Nat Cell Biol. 2004;6(12):1212–20.

24. Spring CM, Kelly KF, O’Kelly I, Graham M, Crawford HC, Daniel JM. The catenin p120ctn inhibits Kaiso-mediated transcriptional repression of the beta-catenin/TCF target gene matrilysin. Experimental cell research. 2005;305(2):253–65.

25. Donaldson NS, Pierre CC, Anstey MI, Robinson SC, Weerawardane SM, Daniel JM. Kaiso represses the cell cycle gene cyclin D1 via sequence-specific and methyl-CpG-dependent mechanisms. PLoS One. 2012;7(11):e50398.

26. Pozner A, Terooatea TW, Buck-Koehntop BA. Cell-specific Kaiso (ZBTB33) Regulation of Cell Cycle through Cyclin D1 and Cyclin E1. J Biol Chem. 2016;291(47):24538–50.

27. Blattler A, Yao L, Wang Y, Ye Z, Jin VX, Farnham PJ. ZBTB33 binds unmethylated regions of the genome associated with actively expressed genes. Epigenetics Chromatin. 2013;6(1):13.

28. Lin QXX, Rebbani K, Jha S, Benoukraf T. ZBTB33 (Kaiso) methylated binding sites are associated with primed heterochromatin. 2019:585653.

29. Moos HK, Patel R, Flaherty SK, Loverde SM, Nikolova EN. H2A.Z facilitates Sox2-nucleosome interaction by promoting DNA and histone H3 tail mobility. Nucleic Acids Res. 2026;54(8).

30. Malaga Gadea FC, Nikolova EN. Structural Plasticity of Pioneer Factor Sox2 and DNA Bendability Modulate Nucleosome Engagement and Sox2-Oct4 Synergism. J Mol Biol. 2023;435(2):167916.

31. Nodelman IM, Patel A, Levendosky RF, Bowman GD. Reconstitution and Purification of Nucleosomes with Recombinant Histones and Purified DNA. Curr Protoc Mol Biol. 2020;133(1):e130.

32. Rueden CT, Schindelin J, Hiner MC, DeZonia BE, Walter AE, Arena ET, et al. ImageJ2: ImageJ for the next generation of scientific image data. BMC Bioinformatics. 2017;18(1):529.

33. Luger K, Rechsteiner TJ, Richmond TJ. Preparation of nucleosome core particle from recombinant histones. Methods Enzymol. 1999;304:3–19.

34. Delaglio F, Grzesiek S, Vuister GW, Zhu G, Pfeifer J, Bax A. NMRPipe: a multidimensional spectral processing system based on UNIX pipes. J Biomol NMR. 1995;6(3):277–93.

35. Raghav SK, Waszak SM, Krier I, Gubelmann C, Isakova A, Mikkelsen TS, et al. Integrative genomics identifies the corepressor SMRT as a gatekeeper of adipogenesis through the transcription factors C/EBPbeta and KAISO. Molecular cell. 2012;46(3):335–50.

36. Hermann A, Goyal R, Jeltsch A. The Dnmt1 DNA-(cytosine-C5)-methyltransferase methylates DNA processively with high preference for hemimethylated target sites. J Biol Chem. 2004;279(46):48350–9.

37. Stutzer A, Liokatis S, Kiesel A, Schwarzer D, Sprangers R, Soding J, et al. Modulations of DNA Contacts by Linker Histones and Post-translational Modifications Determine the Mobility and Modifiability of Nucleosomal H3 Tails. Mol Cell. 2016;61(2):247–59.

38. Morrison EA, Bowerman S, Sylvers KL, Wereszczynski J, Musselman CA. The conformation of the histone H3 tail inhibits association of the BPTF PHD finger with the nucleosome. Elife. 2018;7.

39. Jennings CE, Zoss CJ, Morrison EA. Arginine anchor points govern H3 tail dynamics. Front Mol Biosci. 2023;10:1150400.

40. Cirillo LA, Lin FR, Cuesta I, Friedman D, Jarnik M, Zaret KS. Opening of compacted chromatin by early developmental transcription factors HNF3 (FoxA) and GATA-4. Mol Cell. 2002;9(2):279–89.

41. Zaret KS, Carroll JS. Pioneer transcription factors: establishing competence for gene expression. Genes Dev. 2011;25(21):2227–41.

42. Anderson JD, Widom J. Sequence and position-dependence of the equilibrium accessibility of nucleosomal DNA target sites. J Mol Biol. 2000;296(4):979–87.

43. Li G, Levitus M, Bustamante C, Widom J. Rapid spontaneous accessibility of nucleosomal DNA. Nat Struct Mol Biol. 2005;12(1):46–53.

44. Luo Y, North JA, Rose SD, Poirier MG. Nucleosomes accelerate transcription factor dissociation. Nucleic Acids Res. 2014;42(5):3017–27.

45. Michael AK, Grand RS, Isbel L, Cavadini S, Kozicka Z, Kempf G, et al. Mechanisms of OCT4-SOX2 motif readout on nucleosomes. Science. 2020;368(6498):1460–5.

46. Thapa B, Adhikari NP, Tiwari PB, Chapagain PP. A 5’-Flanking C/G Pair at the Core Region Enhances the Recognition and Binding of Kaiso to Methylated DNA. J Chem Inf Model. 2023;63(7):2095–103.

47. Bostick M, Kim JK, Esteve PO, Clark A, Pradhan S, Jacobsen SE. UHRF1 plays a role in maintaining DNA methylation in mammalian cells. Science. 2007;317(5845):1760–4.

48. Sharif J, Muto M, Takebayashi S, Suetake I, Iwamatsu A, Endo TA, et al. The SRA protein Np95 mediates epigenetic inheritance by recruiting Dnmt1 to methylated DNA. Nature. 2007;450(7171):908–12.

49. Probst AV, Dunleavy E, Almouzni G. Epigenetic inheritance during the cell cycle. Nat Rev Mol Cell Biol. 2009;10(3):192–206.

50. Hashimoto H, Horton JR, Zhang X, Bostick M, Jacobsen SE, Cheng X. The SRA domain of UHRF1 flips 5-methylcytosine out of the DNA helix. Nature. 2008;455(7214):826–9.

51. Avvakumov GV, Walker JR, Xue S, Li Y, Duan S, Bronner C, et al. Structural basis for recognition of hemi-methylated DNA by the SRA domain of human UHRF1. Nature. 2008;455(7214):822–5.

52. Arita K, Ariyoshi M, Tochio H, Nakamura Y, Shirakawa M. Recognition of hemi-methylated DNA by the SRA protein UHRF1 by a base-flipping mechanism. Nature. 2008;455(7214):818–21.

53. Charlton J, Downing TL, Smith ZD, Gu H, Clement K, Pop R, et al. Global delay in nascent strand DNA methylation. Nat Struct Mol Biol. 2018;25(4):327–32.

54. Williams DC, Jr., Cai M, Clore GM. Molecular basis for synergistic transcriptional activation by Oct1 and Sox2 revealed from the solution structure of the 42-kDa Oct1.Sox2.Hoxb1-DNA ternary transcription factor complex. J Biol Chem. 2004;279(2):1449–57.

55. Remenyi A, Lins K, Nissen LJ, Reinbold R, Scholer HR, Wilmanns M. Crystal structure of a POU/HMG/DNA ternary complex suggests differential assembly of Oct4 and Sox2 on two enhancers. Genes Dev. 2003;17(16):2048–59.

56. Shaytan AK, Armeev GA, Goncearenco A, Zhurkin VB, Landsman D, Panchenko AR. Coupling between Histone Conformations and DNA Geometry in Nucleosomes on a Microsecond Timescale: Atomistic Insights into Nucleosome Functions. J Mol Biol. 2016;428(1):221–37.

57. Gatchalian J, Wang X, Ikebe J, Cox KL, Tencer AH, Zhang Y, et al. Accessibility of the histone H3 tail in the nucleosome for binding of paired readers. Nat Commun. 2017;8(1):1489.

58. Strahl BD, Allis CD. The language of covalent histone modifications. Nature. 2000;403(6765):41–5.

59. Kouzarides T. Chromatin modifications and their function. Cell. 2007;128(4):693–705.

60. Schalch T, Duda S, Sargent DF, Richmond TJ. X-ray structure of a tetranucleosome and its implications for the chromatin fibre. Nature. 2005;436(7047):138–41.

61. Sinha D, Shogren-Knaak MA. Role of direct interactions between the histone H4 Tail and the H2A core in long range nucleosome contacts. J Biol Chem. 2010;285(22):16572–81.

62. Allahverdi A, Yang R, Korolev N, Fan Y, Davey CA, Liu CF, et al. The effects of histone H4 tail acetylations on cation-induced chromatin folding and self-association. Nucleic Acids Res. 2011;39(5):1680–91.

63. Ghoneim M, Fuchs HA, Musselman CA. Histone Tail Conformations: A Fuzzy Affair with DNA. Trends Biochem Sci. 2021;46(7):564–78.

64. Tsunaka Y, Furukawa A, Nishimura Y. Histone tail network and modulation in a nucleosome. Curr Opin Struct Biol. 2022;75:102436.

65. Soufi A, Donahue G, Zaret KS. Facilitators and impediments of the pluripotency reprogramming factors’ initial engagement with the genome. Cell. 2012;151(5):994–1004.

66. Buecker C, Srinivasan R, Wu Z, Calo E, Acampora D, Faial T, et al. Reorganization of enhancer patterns in transition from naive to primed pluripotency. Cell Stem Cell. 2014;14(6):838–53.

67. Whyte WA, Bilodeau S, Orlando DA, Hoke HA, Frampton GM, Foster CT, et al. Enhancer decommissioning by LSD1 during embryonic stem cell differentiation. Nature. 2012;482(7384):221–5.

68. Hu S, Wan J, Su Y, Song Q, Zeng Y, Nguyen HN, et al. DNA methylation presents distinct binding sites for human transcription factors. Elife. 2013;2:e00726.

69. Lee JE, Wang C, Xu S, Cho YW, Wang L, Feng X, et al. H3K4 mono- and di-methyltransferase MLL4 is required for enhancer activation during cell differentiation. Elife. 2013;2:e01503.

70. Local A, Huang H, Albuquerque CP, Singh N, Lee AY, Wang W, et al. Identification of H3K4me1-associated proteins at mammalian enhancers. Nat Genet. 2018;50(1):73–82.

71. Tsunaka Y, Ohtomo H, Morikawa K, Nishimura Y. Partial Replacement of Nucleosomal DNA with Human FACT Induces Dynamic Exposure and Acetylation of Histone H3 N-Terminal Tails. iScience. 2020;23(10):101641.

72. Bohne F, Langer D, Martine U, Eider CS, Cencic R, Begemann M, et al. Kaiso mediates human ICR1 methylation maintenance and H19 transcriptional fine regulation. Clin Epigenetics. 2016;8:47.

73. Polach KJ, Widom J. Mechanism of protein access to specific DNA sequences in chromatin: a dynamic equilibrium model for gene regulation. J Mol Biol. 1995;254(2):130–49.

