## Supplemental Data for "Kaiso reads methylated CpGs at nucleosome entry/exit and displaces the H3 tail"

**Table S1.** Nucleotide sequences for Widom 601 substrates modified with the methyl-CpG Kaiso binding site ($MeCG_2$, underlined). The dyad ($0^{th}$) position is bolded.

| |
|---|
| **Widom 601 (145 base pair)**<br>(NCP 601)<br>5’-TG GAGAATCCCG GTGCCGAGGC CGCTCAATTG GTCGTAGACA GCTCTAGCAC CGCTTAAACG CACGTACGCG **C** TGTCCCCCGC GTTTTAACCG CCAAGGGGAT TACTCCCTAG TCTCCAGGCA CGTGTCAGAT ATATACATCC TG-3’ |
| **Modified 601 sequences (5’FAM-labeled strand; 147 base pair)**<br>(NCP SHL 6.5)<br>5'-TG GAGAATCCCG GTGCCGAGGC CGCTCAATTG GTCGTAGACA GCTCTAGCAC CGCTTAAACG CACGTACGCG **C** TGTCCCCCGC GTTTTAACCG CCAAGGGGAT TACTCCCTAG TCTCCAGGCA CGTGTCAGAT TCTCGCGAGA TGTC-3’<br>(NCP SHL 5.5)<br>5'-TG GAGAATCCCG GTGCCGAGGC CGCTCAATTG GTCGTAGACA GCTCTAGCAC CGCTTAAACG CACGTACGCG **C** TGTCCCCCGC GTTTTAACCG CCAAGGGGAT TACTCCCTAG TCTCCAGGCA TCTCGCGAGA ATATACATCC TGTC-3’<br>(NCP SHL 2.5)<br>5'-TG GAGAATCCCG GTGCCGAGGC CGCTCAATTG GTCGTAGACA GCTCTAGCAC CGCTTAAACG CACGTACGCG **C** TGTCCCCCGC GTTTTAACCG TCTCGCGAGA TACTCCCTAG TCTCCAGGCA CGTGTCAGAT ATATACATCC TGTC-3’<br>(NCP SHL 0.5)<br>5’-TG GAGAATCCCG GTGCCGAGGC CGCTCAATTG GTCGTAGACA GCTCTAGCAC CGCTTAAACG CACGTACGCG **C** TCTCGCGAGA GTTTTAACCG CCAAGGGGAT TACTCCCTAG TCTCCAGGCA CGTGTCAGAT ATATACATCC TGTC-3’ |

**Table S2.** Cy3-labeled primer sequences used for PCR of modified 601 substrates in Table S1. X = 5-methylcytosine in hemi-methylated substrates; X = cytosine in unmethylated substrates.

| |
|---|
| (NCP SHL 6.5)<br>5'-GACATCTXGXGAGAATCTGACACGTGCCTGG-3'<br>(NCP SHL 5.5)<br>5'-GACAGGATGTATATTCTXGXGAGATGCCTGG-3’<br>(NCP SHL 2.5)<br>5'-GACAGGATGTATATATCTGACACGTGCCTGGAGACTAGGGAGTATCTXGXG-3’<br>(NCP SHL 0.5)<br>5’-GACAGGATGTATATATCTGACACGTGCCTGGAGACTAGGGAGTAATCCCCTTGGCGGTTAAAACTCTXGXG-3’ |

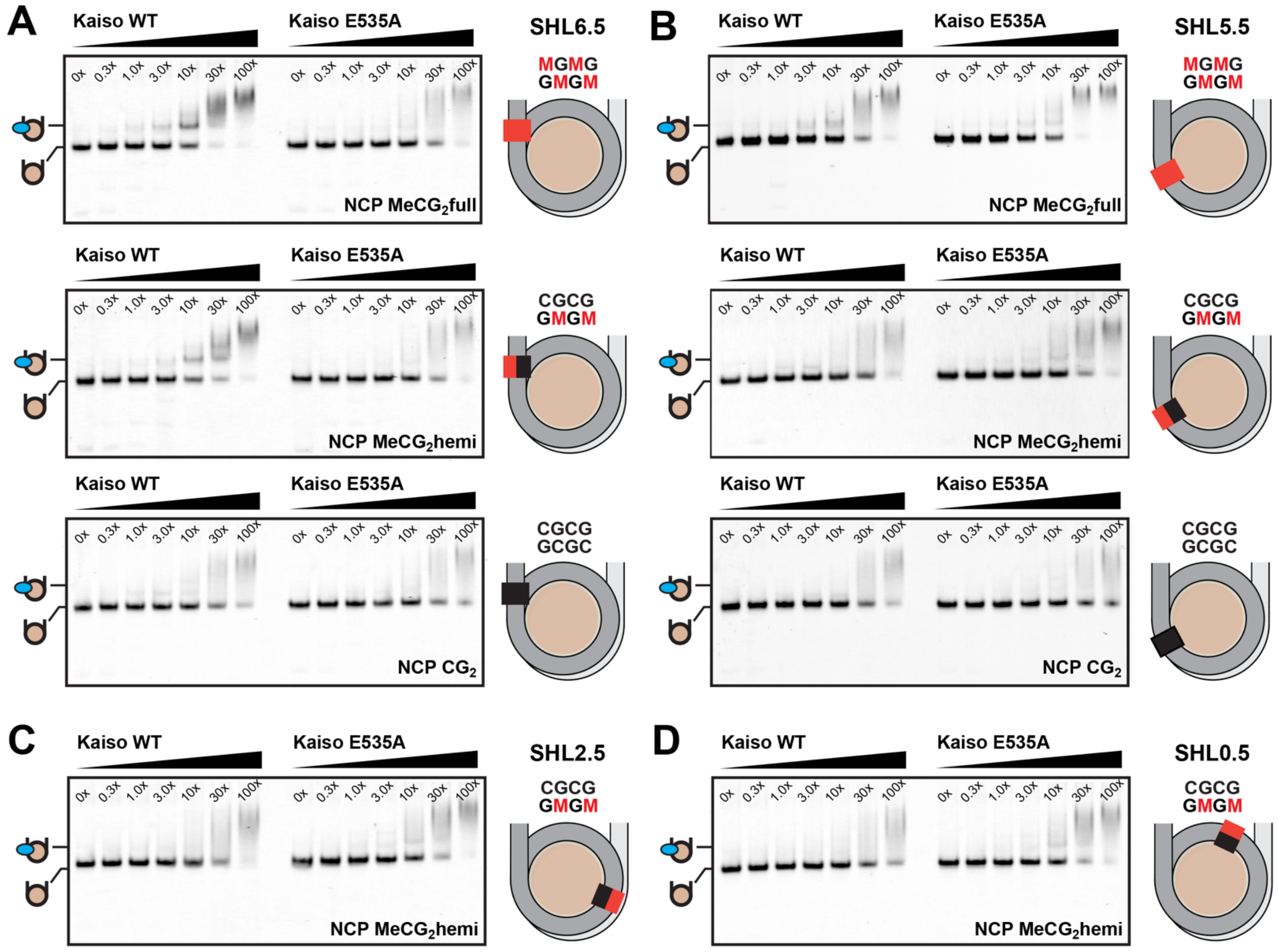


**Figure S1. Kaiso binding to methylated nucleosomes is position- and methylation-dependent.** Representative EMSA gels of WT and E535A Kaiso (0, 0.3, 1, 3, 10, 30, and 100X) titrated against 10 nM nucleosomes (NCPs) containing Kaiso binding site at **(A)** SHL 6.5, **(B)** SHL 5.5, **(C)** SHL 2.5, **(D)** SHL 0.5. Schematics on the right indicate the methylation state of the central CpG steps at the Kaiso site in each substrate, including fully methylated (NCP $MeCG_2$full), hemi-methylated (NCP $MeCG_2$hemi), and unmethylated (NCP $CG_2$) nucleosomes.

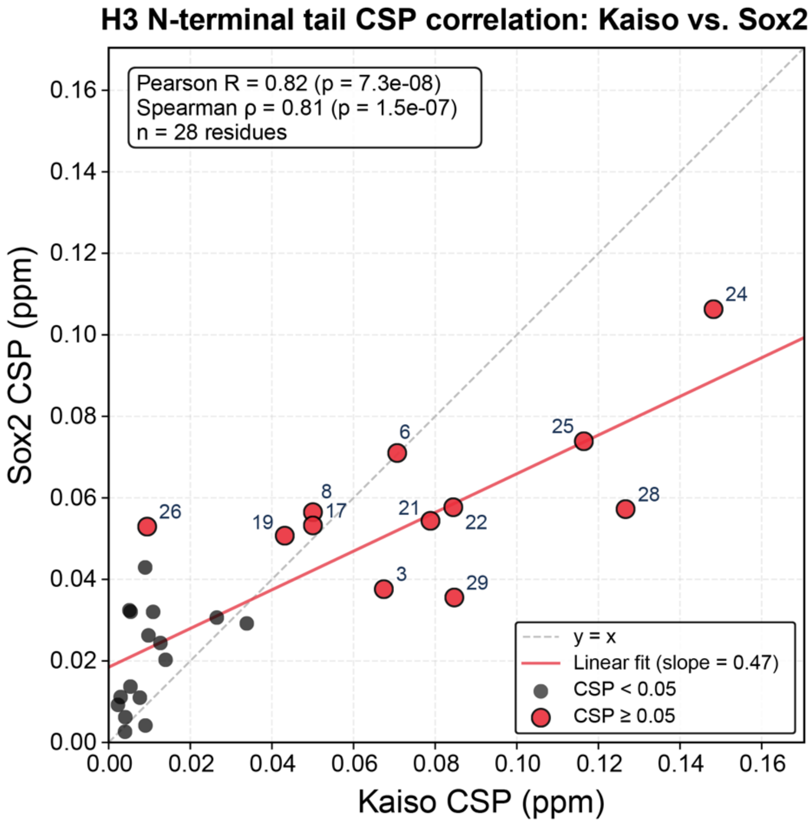


**Figure S2. Kaiso and Sox2 perturb a near-identical set of H3 N-terminal tail residues.** Per-residue chemical shift perturbations (CSPs) for $^{15}$N-labeled H3 in Kaiso-bound NCP MeCG$_2$hemi (SHL 6.5) are plotted against CSPs from previously published Sox2-bound NCP 62$_F$ nucleosomes.(1) Each point represents one H3 tail residue (n = 28), with hotspots (CSP ≥ 0.05 ppm in either dataset) shown as filled circles and labeled. Dashed line, y = x; red line, linear best fit (slope = 0.47). The fingerprints are strongly correlated (Pearson R = 0.82, $p = 7 \times 10^{-8}$; Spearman ρ = 0.81), despite the two proteins recognizing nucleosomal DNA through fundamentally different chemistries.

### References

1. Moos HK, Patel R, Flaherty SK, Loverde SM, Nikolova EN. H2A.Z facilitates Sox2-nucleosome interaction by promoting DNA and histone H3 tail mobility. Nucleic Acids Res. 2026;54(8).
